## Supplementary information for "Coupled equilibria of dimerization and lipid binding modulate SARS Cov 2 Orf9b interactions and interferon response"

### Supplemental Figures

#### Table 1:

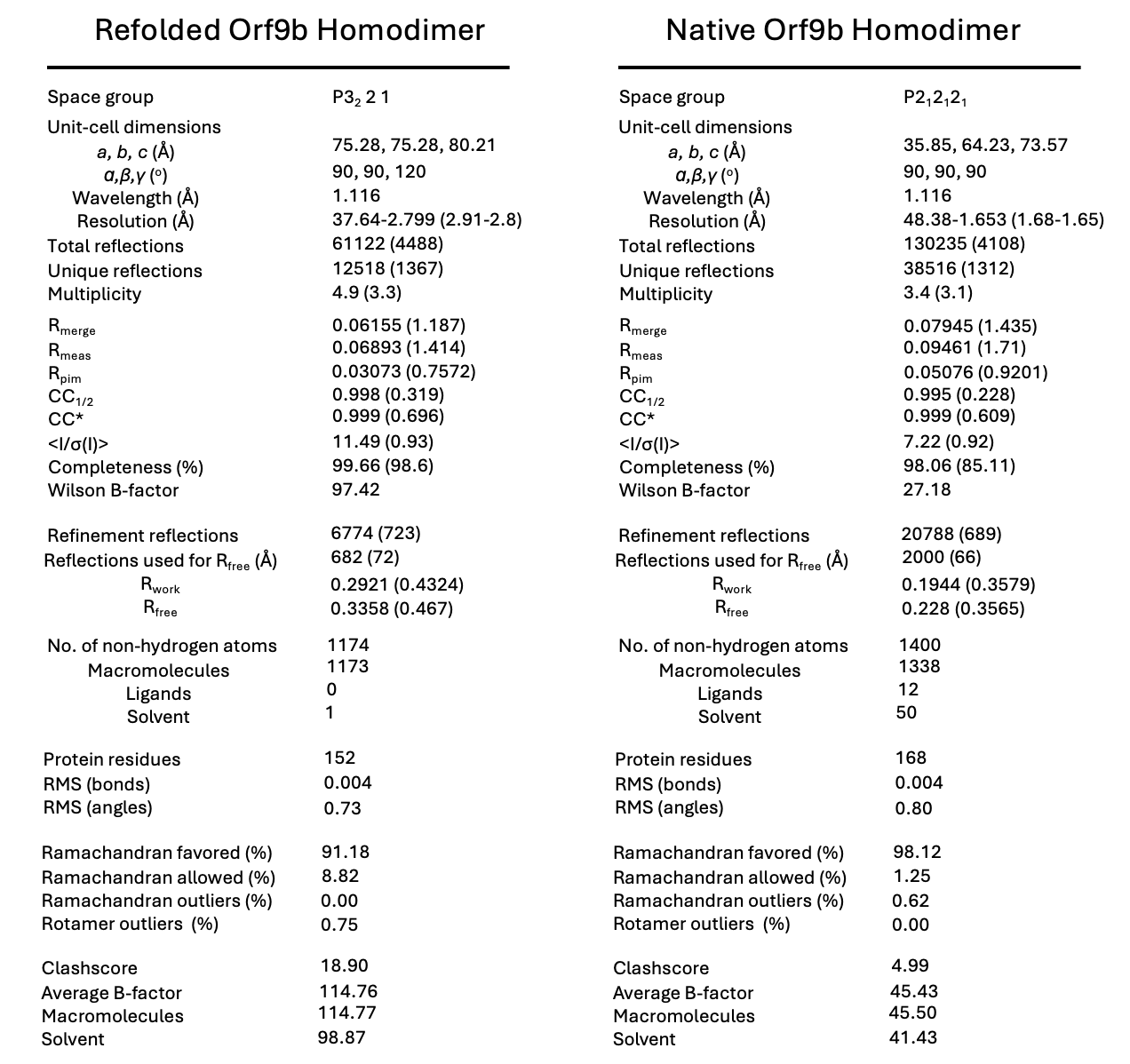

#### Figure 1 - Supplemental 1:

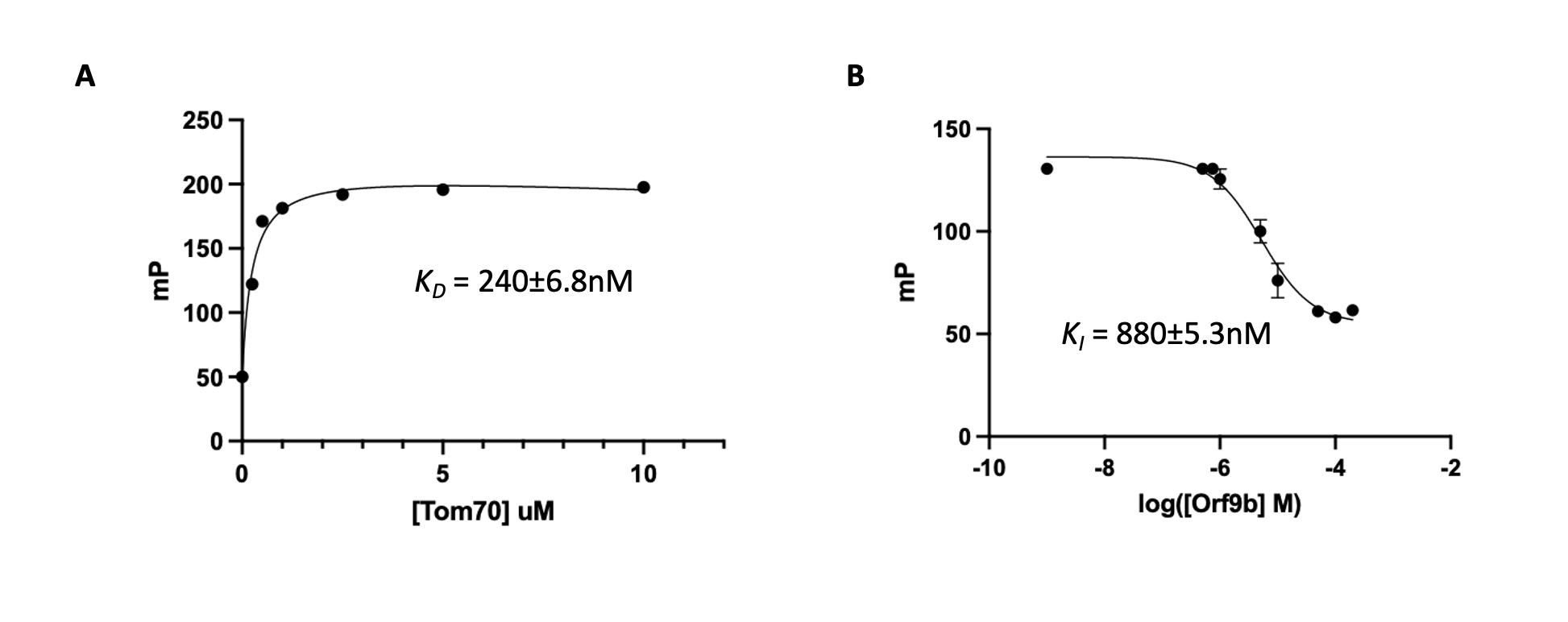

A. Determining the K_D_ of Orf9b-FITC:Tom70 by titration of Tom70 against a fixed concentration of Orf9b-FITC. 200nM of Orf9b-FITC was kept constant and titrated with increasing concentrations of Tom70. A non-linear regression model using 1:1 binding was used to calculate the K_D_. The K_D_ was 240±6.8nM
B. Determining the K_i_ of unlabeled Orf9b peptide by competition binding. Orf9b peptide was titrated against 200nM Orf9b-FITC and 2.5uM of Tom70. Non-linear regression using a one-site competitive binding model was used to calculate the K_i_ of the unlabeled Orf9b peptide using the calculated K_D_ of Orf9b-FITC:Tom70 from (A). The unlabeled Orf9b peptide K_i_ was 880±5.3nM

#### Figure 1 - Supplemental 2:

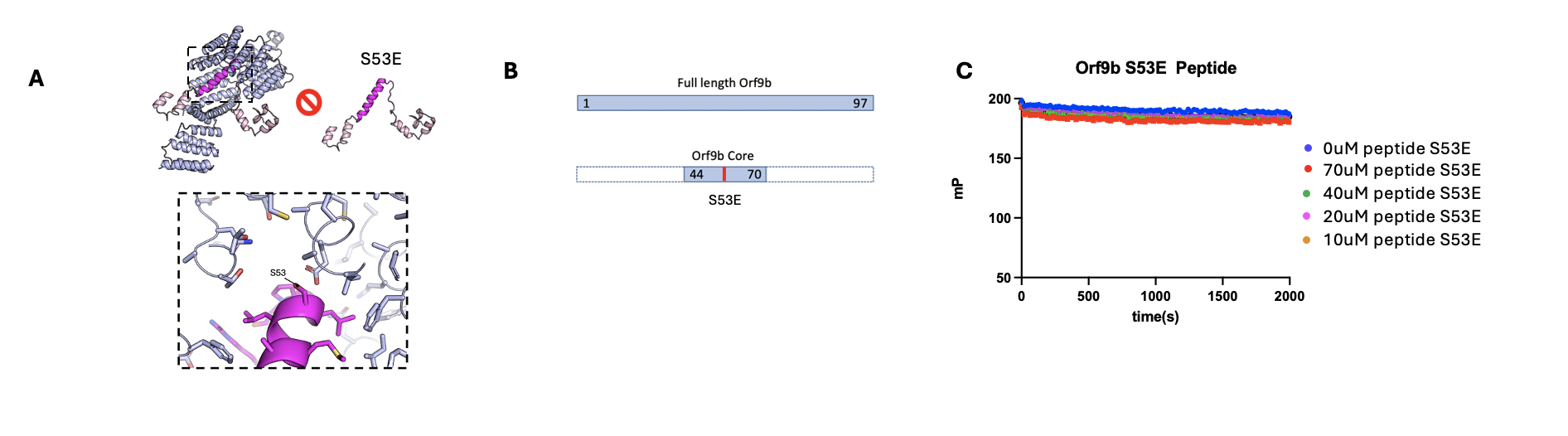

A. Phosphorylated Orf9b at S53 does not bind to Tom70. Zoomed in view of Orf9b bound to Tom70 showing the location of Orf9b S53 relative to the surrounding Tom70 residues.
B. Illustration of the Orf9b S53E peptide derived from the full length WT Orf9b sequence.
C. Orf9b S53E peptide FP kinetic assay results showing that the phosphomimetic peptide does not bind to Tom70.

#### Figure 1 - Supplemental 3:

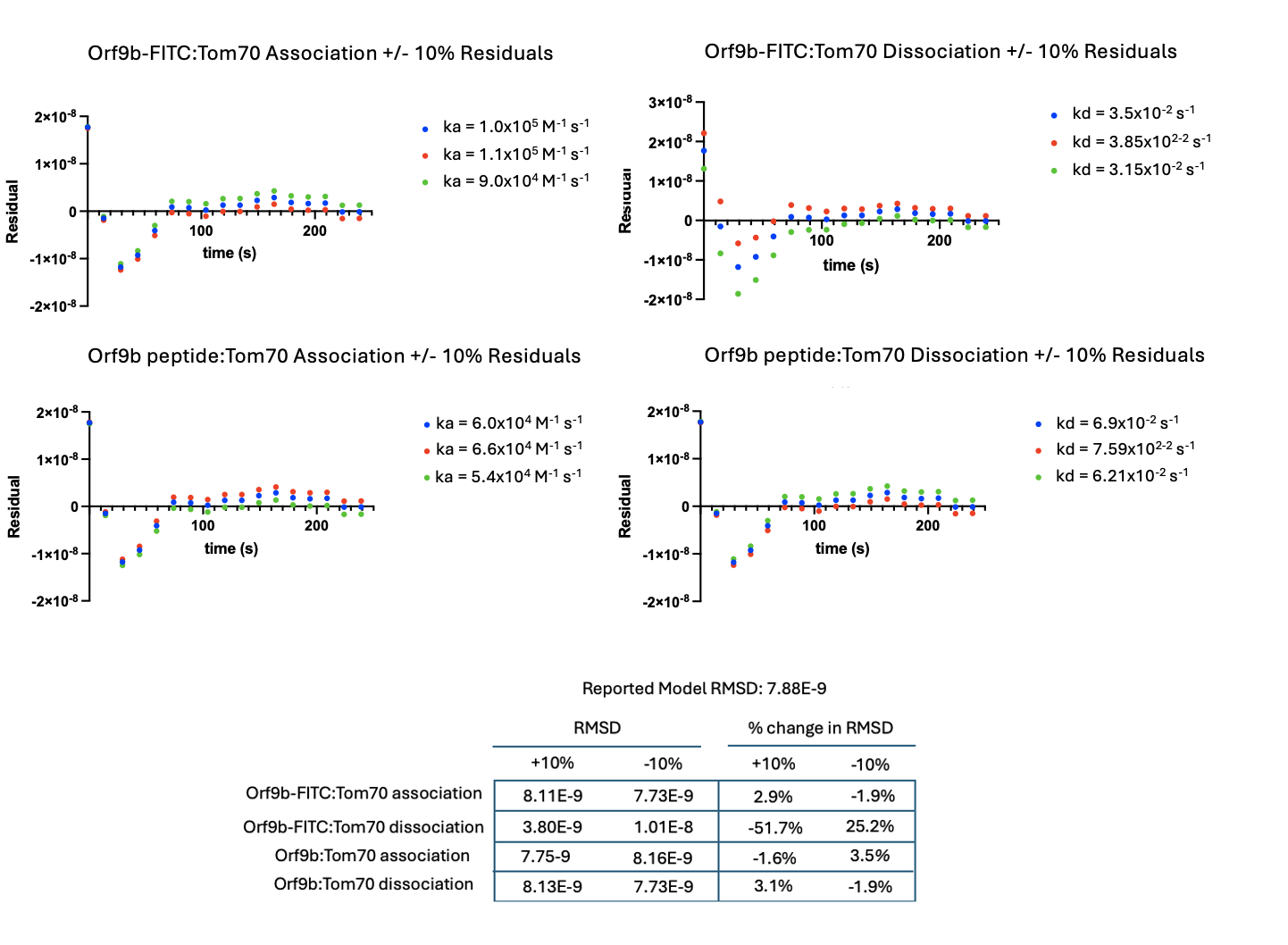

Plots of residuals from Orf9b peptide model showing effect of an increase or decrease by 10% on each model parameter. All residuals and reporting are with respect to the100uM of unlabeled Orf9b peptide condition. Blue dots: reported value. Red dots: 10% increase in reported value. Green dots: 10% decrease in reported value. Table reporting of RMSD values for model fitsafter +/-10% change to model parameter (Left column) and percent change in RMSD relative to reported model RMSD (Right column).

#### Figure 2 - Supplemental 1:

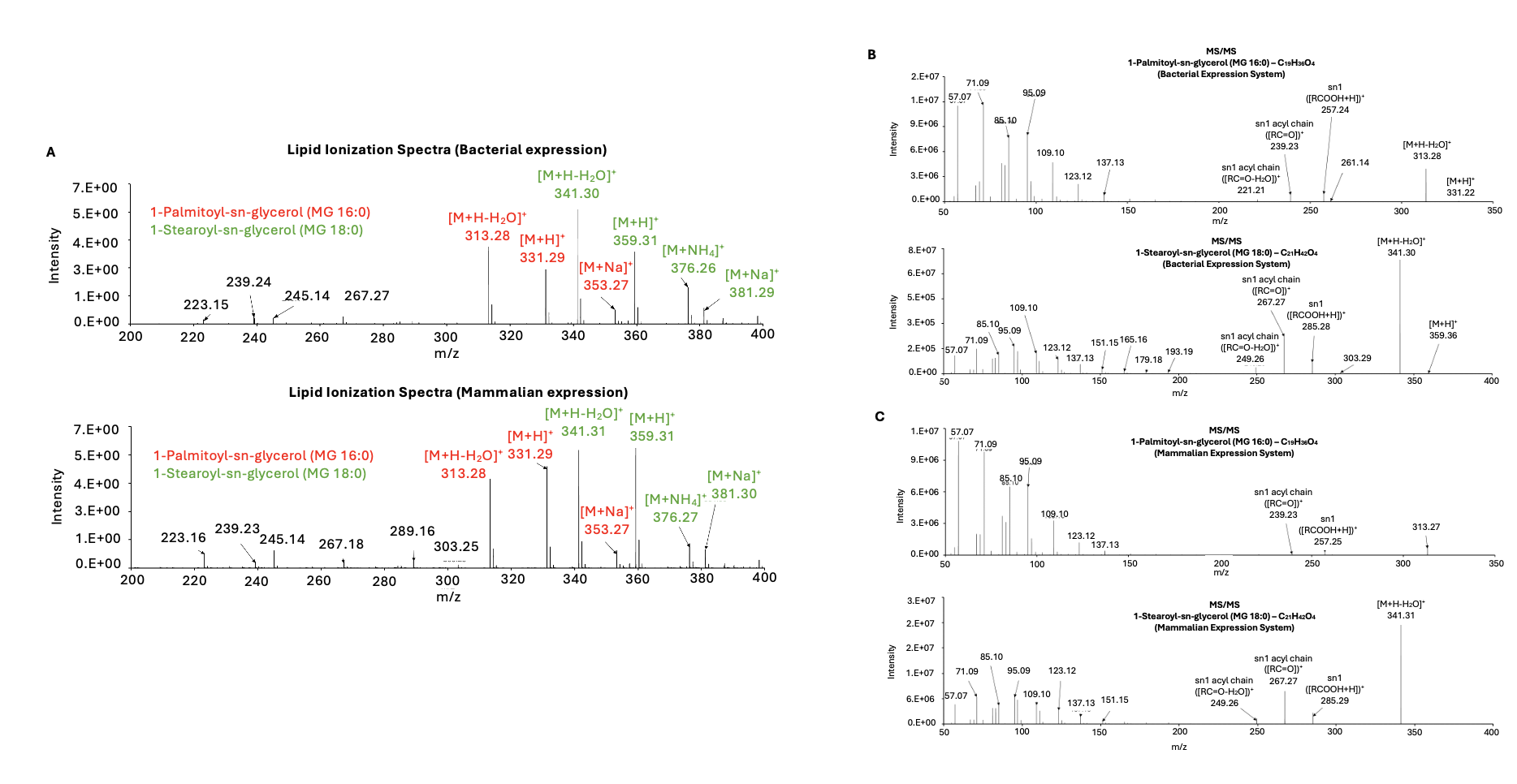

A. Mass spectra of lipids extracted from natively folded Orf9b in both (top) E.coli and (bottom) mammalian expression systems.
B. (Top) Tandem mass spectrometry (MS/MS) ionization spectra of 1-Palmitoyl-sn-glycerol and (bottom)1-Strearoyl-sn-glycerol from bacterial expression systems.
C. (Top) Tandem mass spectrometry (MS/MS) ionization spectra of 1-Palmitoyl-sn-glycerol and (bottom) 1-Strearoyl-sn-glycerol from mammalian expression systems.

#### Figure 2- Supplemental 2:

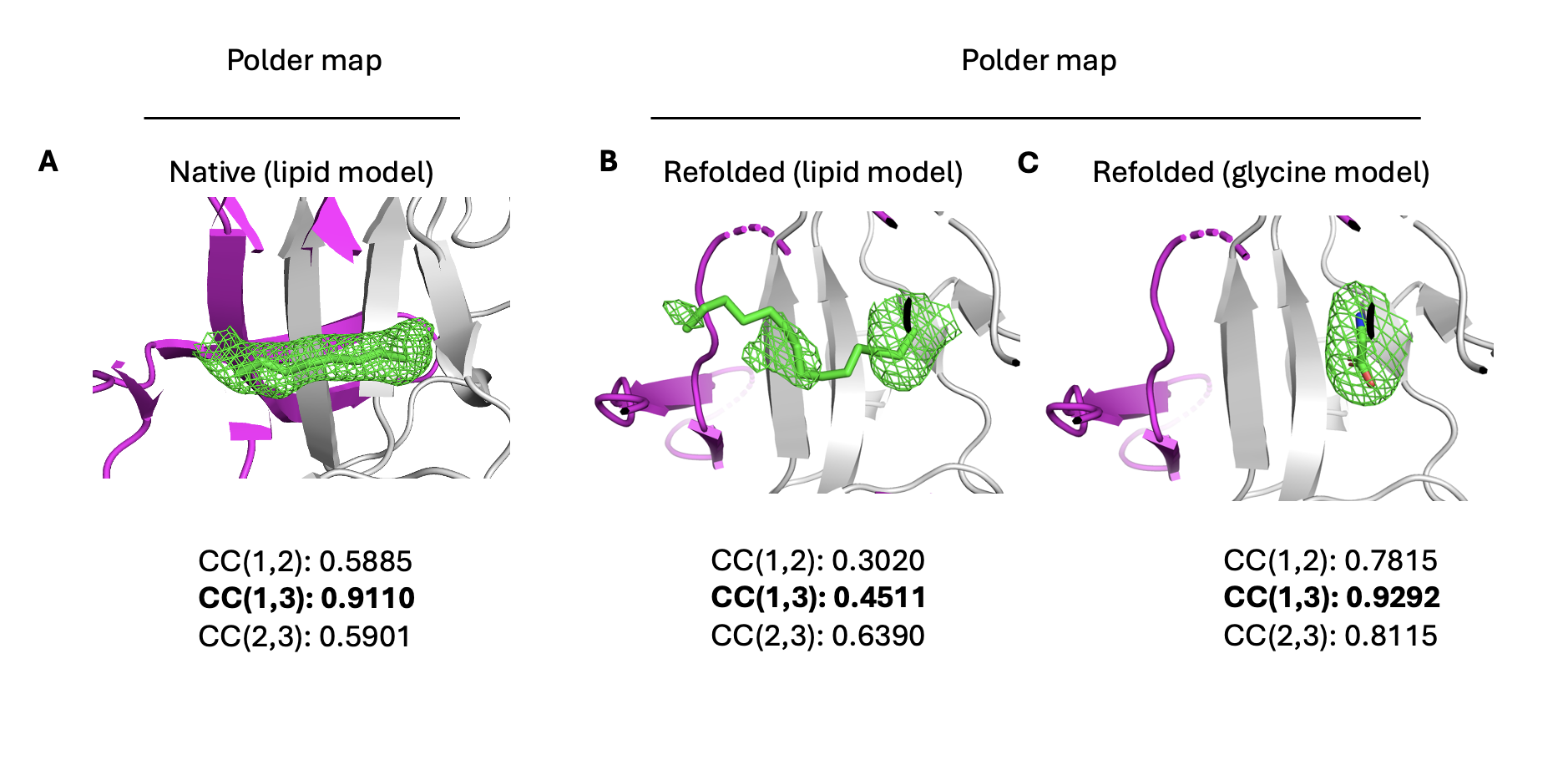

A. Polder map calculated for the lipid-molecule bound to the Orf9b homodimer. Polder map shows strong support for the presence of the lipid with CC1,3 value being 0.9110.
B. Polder map calculated for the refolded Orf9b homodimer with the lipid molecule modeled into the central channel. Polder maps do not support the placement of the lipid with CC1,3 value being 0.4511 indicating that residual Fo-Fc and 2Fo-Fc density peaks are explained by either noise or the bulk solvent mask.
C. Polder maps calculated for the refolded Orf9b homodimer with glycine (from crystallization conditions) modeled into the largest 2Fo-Fc peak within the central channel. Polder maps plausibly support placement of glycine within the central channel with CC1,3 value being 0.9292.

#### Figure 3 - Supplemental 1

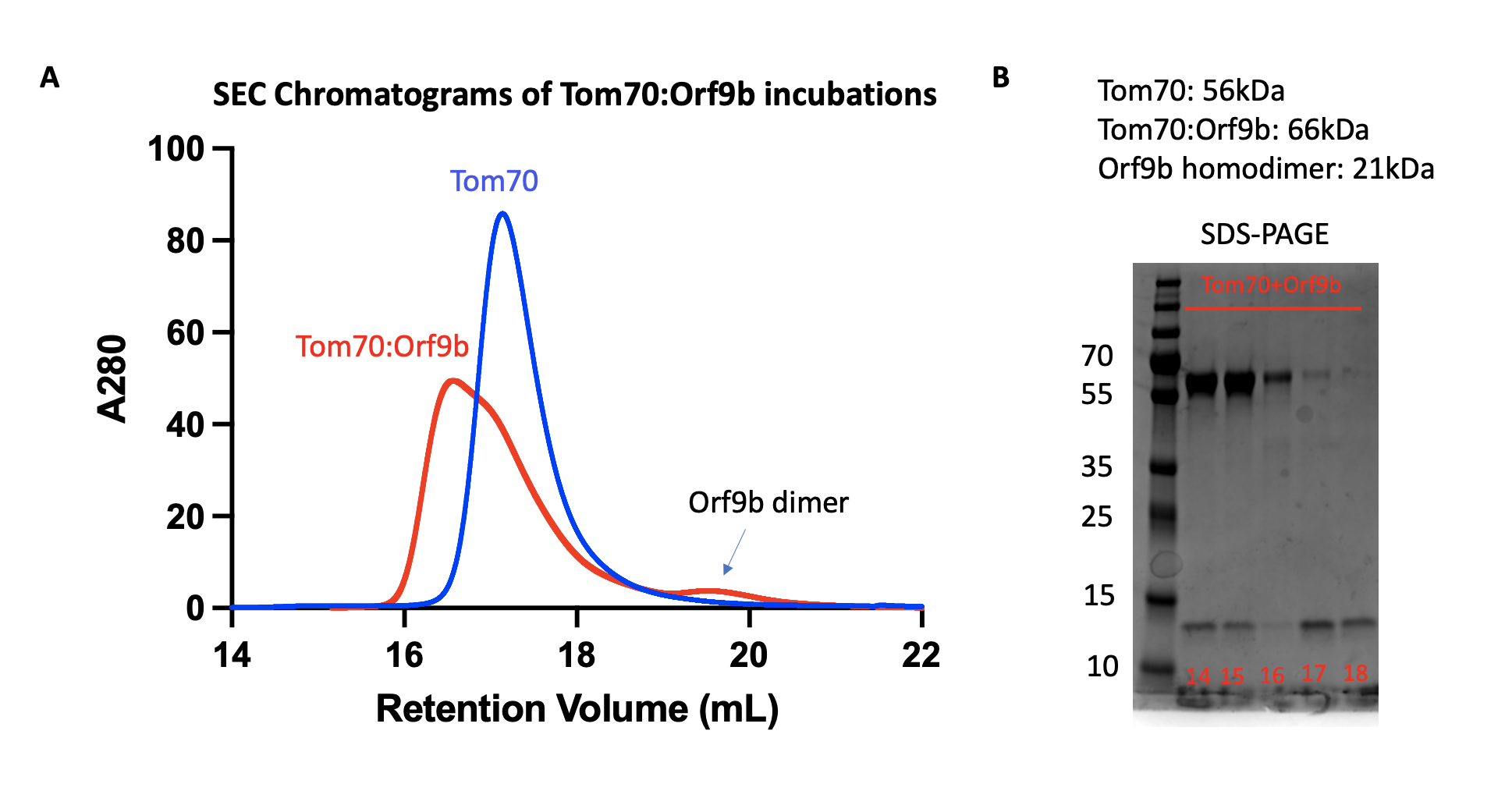

A. Size exclusion chromatogram of Tom70:Orf9b, Tom70, and Orf9b overlaid. Refolded Orf9b and Tom70 show a left-shift relative to Tom70 alone indicating the formation of a higher molecular weight complex.
B. SDS-polyacrylamide gel of the Tom70:Orf9b SEC fractions with molecular weights of all possible species present. The first fractions (14-15) show Tom70 and Orf9b eluting together which is indicative of the Tom70:Orf9b complex followed by a fraction that is mostly Tom70 (16) and then two fractions containing mostly Orf9b (17-18).

#### Figure 4 - Supplemental 1:

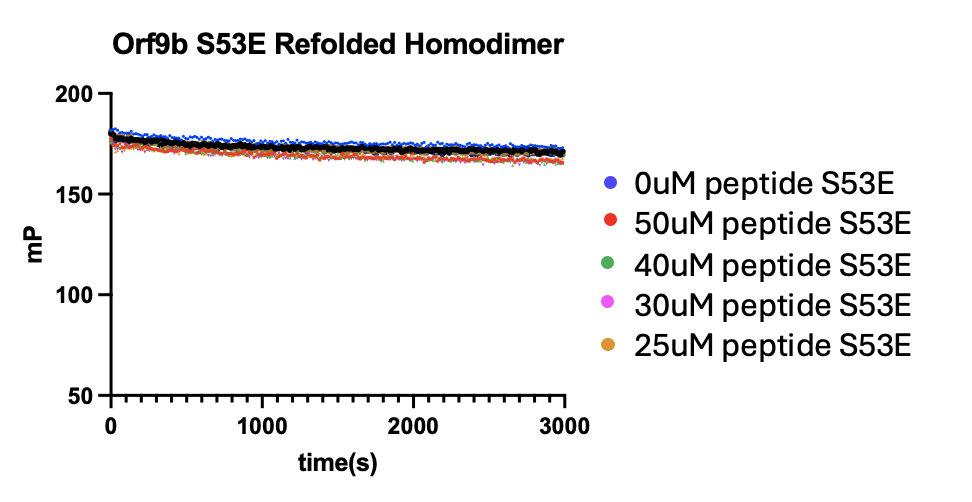

Orf9b S53E Refolded homodimer FP kinetic assay results showing that the phosphomimetic mutation does not bind to Tom70.

#### Figure 4 - Supplemental 2:

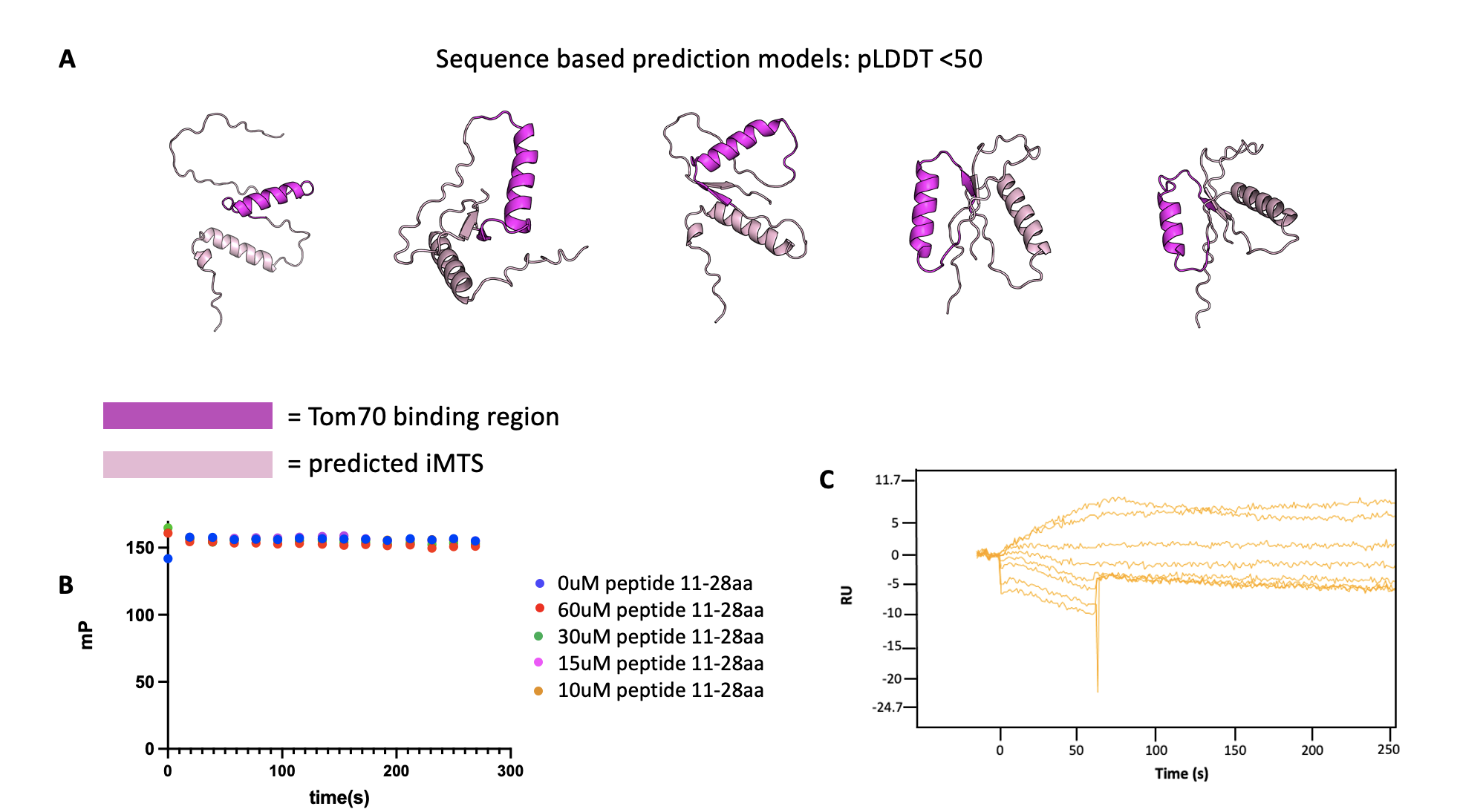

A. AlphaFold predictions of monomeric Orf9b without structural templates used. The region of Orf9b that is known to bind to Tom70 is highlighted in magenta and the residues that form the Orf9b homodimer interface are highlighted in pink. A second helix is predicted to form in monomeric Orf9b comprising residues 11-28. The top 5 predicted structures are shown with pLDDT scores less than 50 indicating a low confidence in the predicted monomeric structure of Orf9b.
B. FP kinetic assay using a peptide composed of residues 11-28 shows no binding to Tom70 at the C-terminal domain where the structurally resolved portion of Orf9b binds.
C. SPR using a peptide composed of residues 11-28 against immobilized Tom70 shows no binding indicating that the second predicted helix does not bind to Tom70.

#### Figure 4 - Supplemental 3:

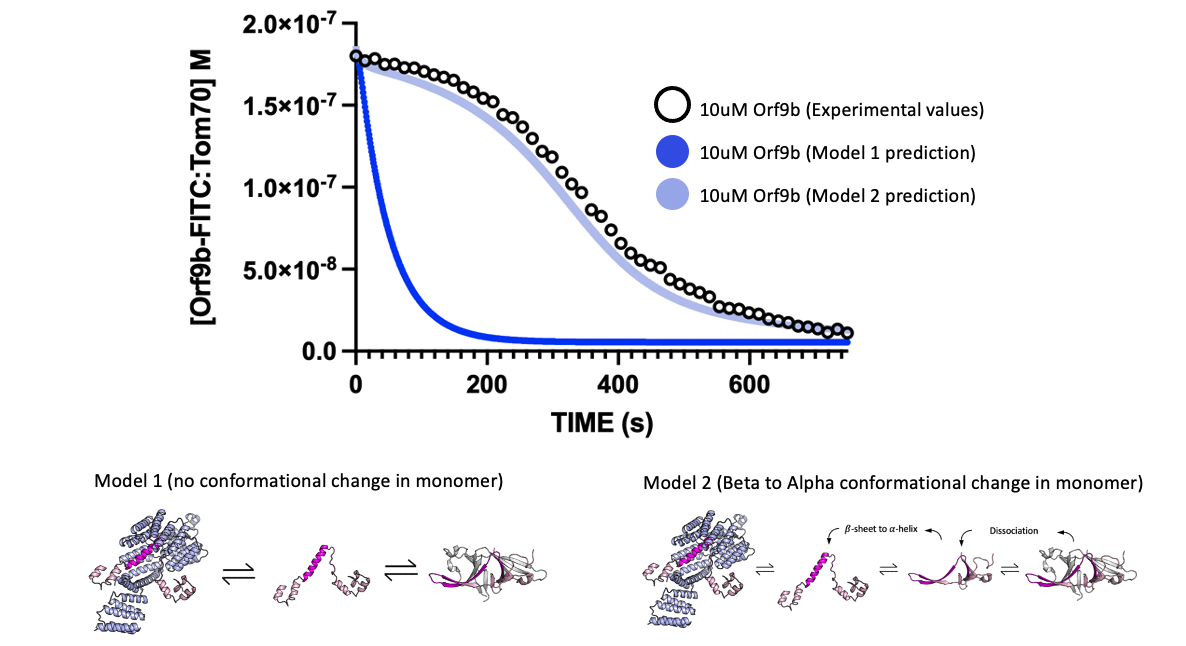

Comparison of kinetic model 1 and 2 in describing experimental results from the kinetic binding assay. Experimental results using 10uM of refolded Orf9b homodimer are shown as rings with the predicted behavior of model 1 (equilibrium exchange) shown as a dark blue line. The predicted behavior of model 2 (equilibrium exchange with a conformational change between β-sheet and ɑ-helical monomers) is shown as the light blue line. Model parameter values were the same as described in Figure 4D and kept constant in both model comparisons.

#### Figure 4 - Supplemental 4:

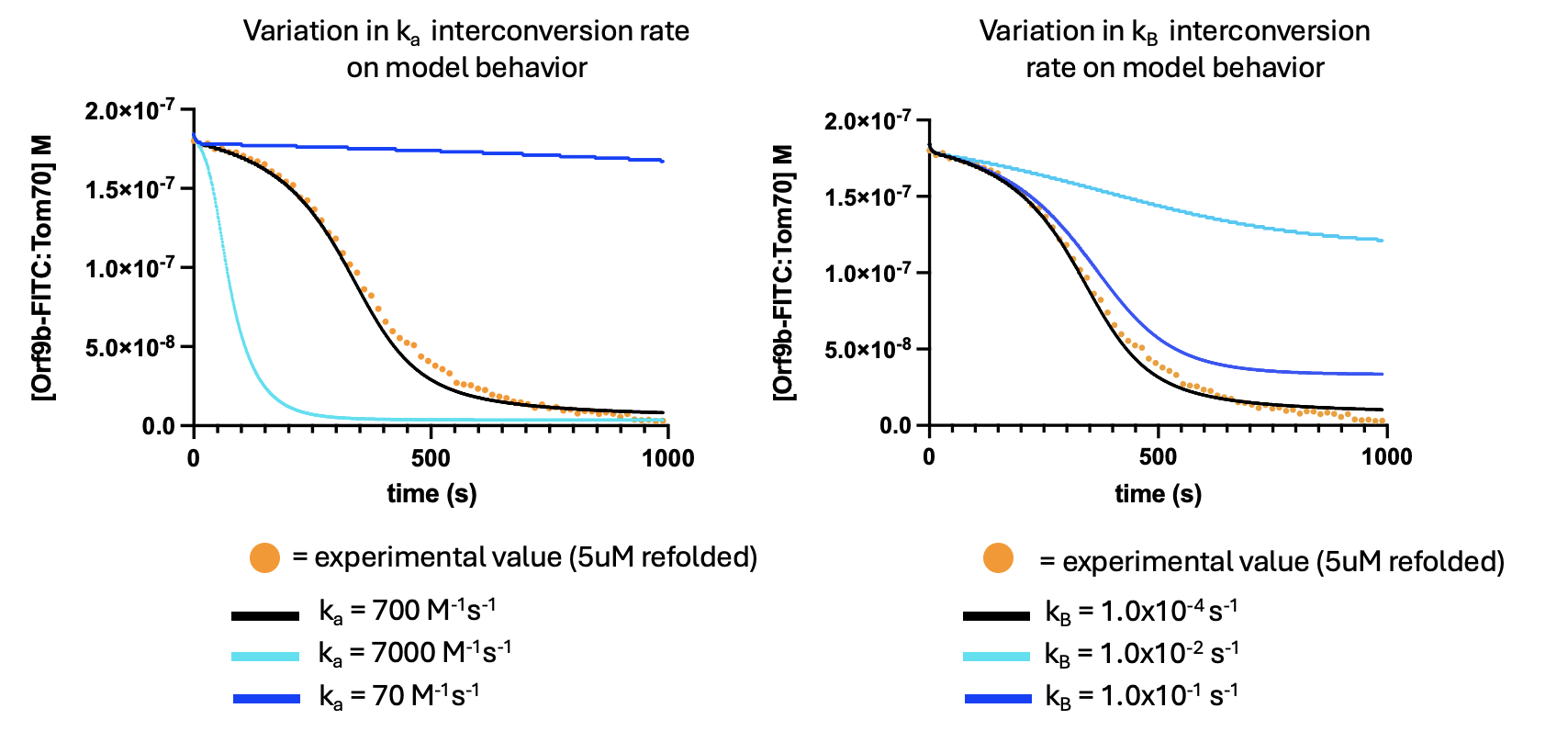

Plots of model behavior showing the effect of changes to alpha-beta and beta-alpha monomer interconversion rates compared to experimental values. Data is modeled with respect to the apo-Orf9b homodimer 5uM condition. Black line represents reported model fit and values used.

#### Figure 4 - Supplemental 5:

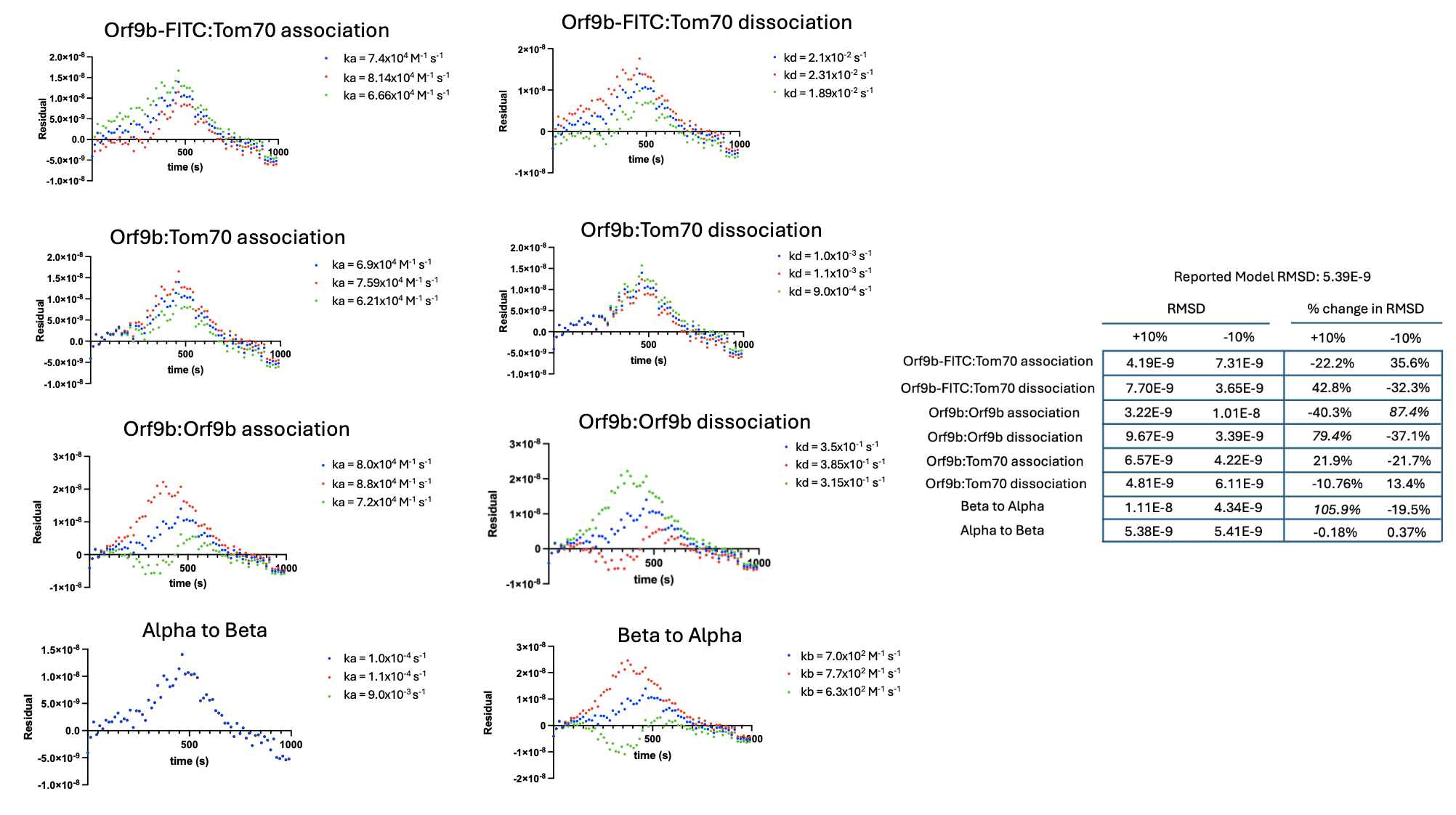

Plot of residuals showing the effect of increasing or decreasing individual model parameters 10% compared to the reported values. All residual plots are with respect to the 5uM apo-Orf9b homodimer condition. Blue dots: reported value. Red dot: 10% increase in reported value. Green dot: 10% decrease in reported value. (Left columns) Table of RMSD values calculated from model fits showing the effect of both +/-10% change to individual model parameters. (Right columns) Percent change in RMSD values subjected to +/-10% change for individual model parameters relative to the RMSD of the reported model.

#### Figure 5 - Supplemental 1

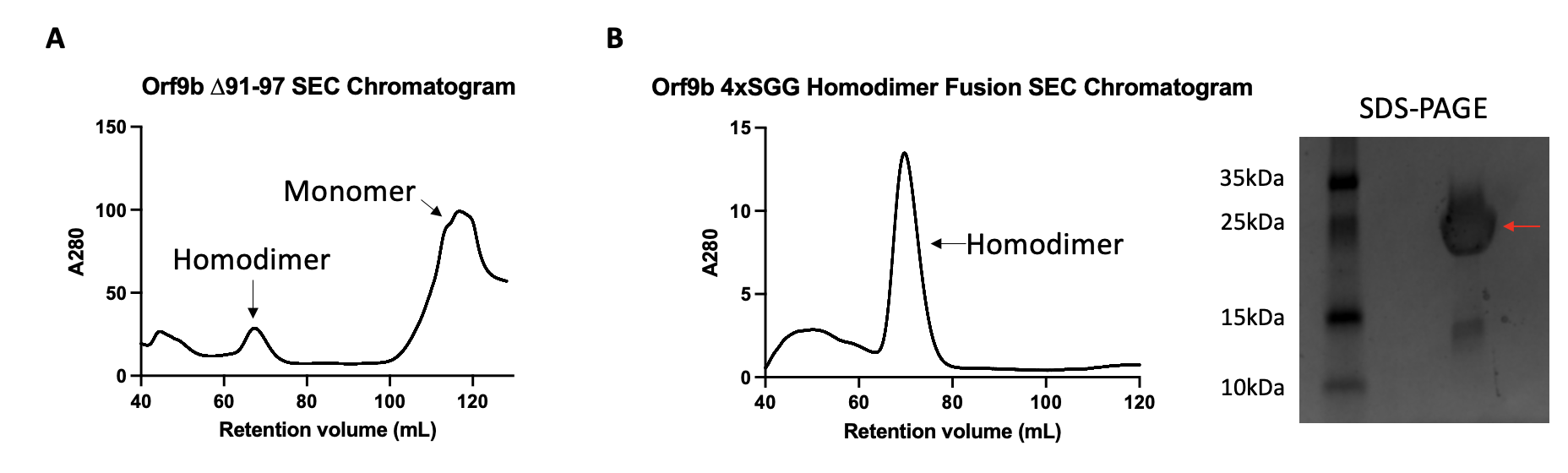

A. Size exclusion chromatography chromatogram of the Orf9b ∆91-97 truncated construct. The predicted homodimer has a peak centered at a retention volume of 70mL followed by a much larger peak that eluted at the end of the column volume which we attribute to the monomeric species.
B. (Left) Size exclusion chromatography chromatogram of the Orf9b 4xSGG fusion construct. The predicted homodimer has a retention volume of 70mL. (Right) SDS-PAGE of the homodimer elution peak showing the predicted molecular weight of the Orf9b fusion construct of ~25kDa (red arrow). The minor band between 10kDa and 15kDa we attribute to a minor degradation product.

#### Figure 6 - Supplemental 1

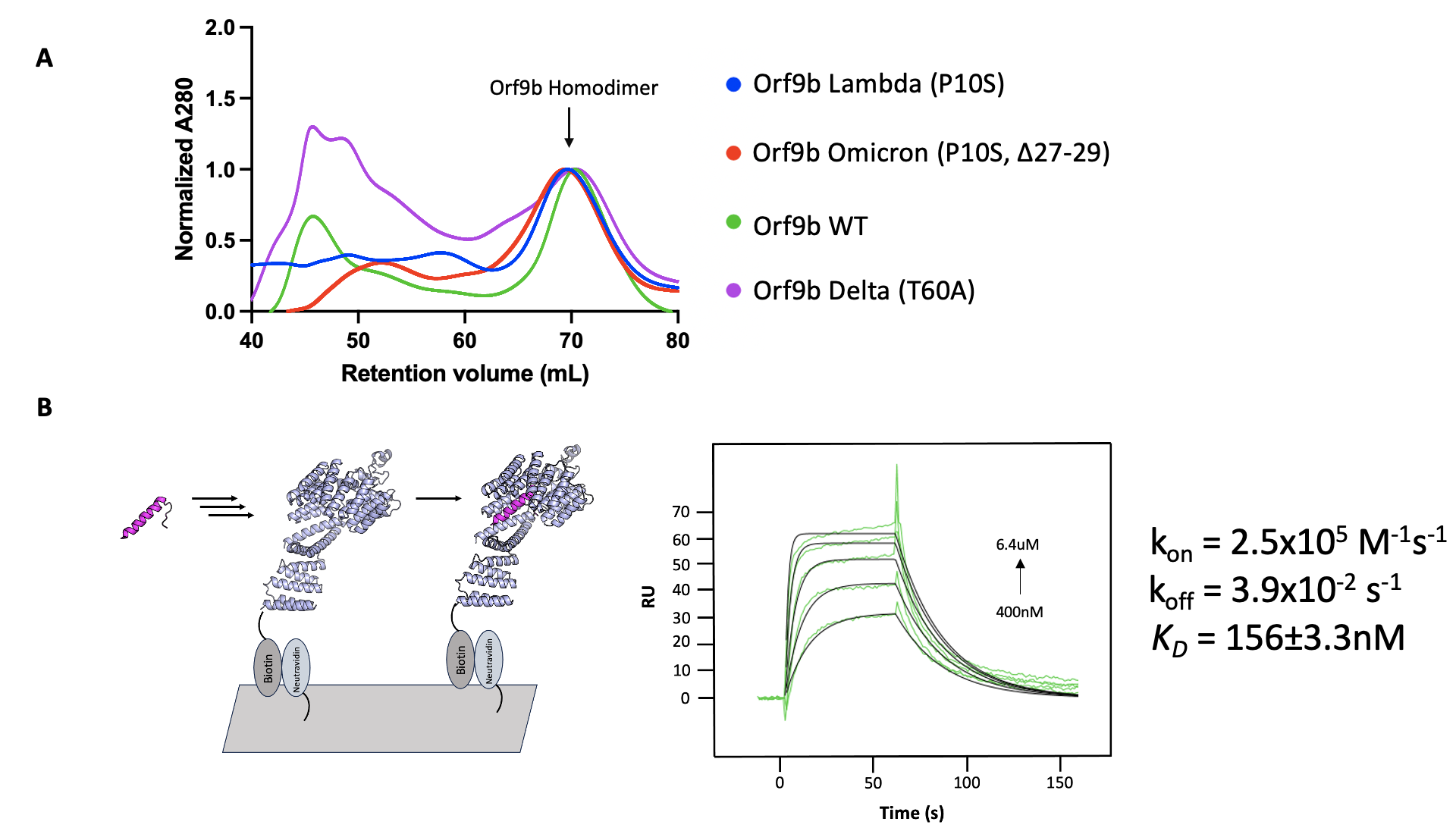

A. Size exclusion chromatogram showing overlaid chromatograms of refolded WT, Delta, Lambda, and Omicron Orf9b homodimers with normalized A280 absorbance. The retention volumes corresponding to the homodimer are centered at ~70mL in all cases indicating recovery of the homodimer after refolding.
B. (Left) SPR diagram of immobilized Tom70 and Orf9b peptide used as an analyte. (Right) SPR response curves using Orf9b T60A peptide as the analyte. Experimental response curves are shown in green and global 1:1 binding fits are shown in black with corresponding kinetic rate constants and K_D_.

#### Figure 6 - Supplemental 2

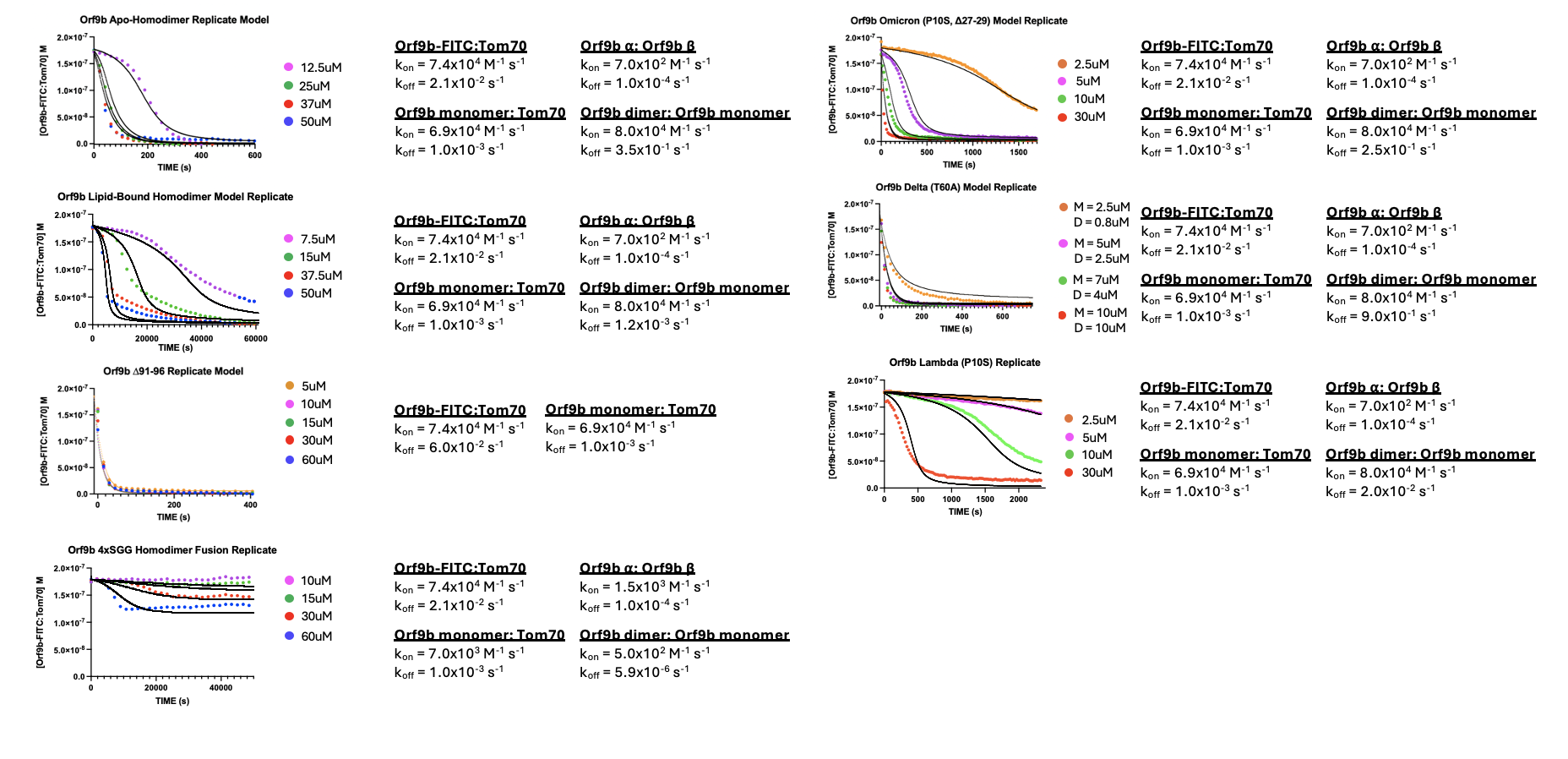

Biological replicates of all Orf9b variants and constructs performed in FP kinetic assay with model results (solid black lines) and parameters used. All concentrations refer to the Orf9b homodimer concentrations unless specified otherwise (see Delta T60A).
